# Structure of the N-RNA/P interface reveals mode of L/P attachment to the nucleocapsid of human metapneumovirus

**DOI:** 10.1101/2023.06.10.544460

**Authors:** Jack D. Whitehead, Hortense Decool, Cédric Leyrat, Loic Carrique, Jenna Fix, Jean-François Eléouët, Marie Galloux, Max Renner

## Abstract

Human metapneumovirus (HMPV) is a major cause of respiratory illness in young children. The polymerase complex of HMPV consists of two obligate components, the L polymerase and its cofactor, the phosphoprotein P. During replication and transcription, the L/P complex traverses the viral RNA genome, which is encapsidated within multimerized N nucleoproteins. An essential interaction between N and a C-terminal region of P is required for tethering of the L/P polymerase to the RNA template. This N-P interaction is also involved in the formation of cytoplasmic viral factories in infected cells, called inclusion bodies. To define how L/P recognizes N-encapsidated RNA (N-RNA) we employed cryogenic electron microscopy (cryo-EM) and molecular dynamics simulations, coupled to polymerase activity assays and imaging of inclusion bodies in transfected cells. We report a 2.9 Å resolution structure of a triple-complex between multimeric N, bound to both RNA and the C-terminal region of P. Furthermore, we also present cryo-EM structures of assembled N in different oligomeric states, highlighting the plasticity of N. Combined with our functional assays, these structural data delineate in molecular detail how P attaches to N-RNA whilst retaining substantial conformational dynamics. Moreover, the N-RNA-P triple complex structure provides a molecular blueprint for the design of therapeutics to potentially disrupt the attachment of L/P to its template.

---

Human Metapneumovirus (HMPV) was first reported in 2001 when it was isolated from children in the Netherlands with symptoms ranging from mild respiratory disease to severe pneumonia^1^. HMPV is thought to have a zoonotic origin following a spill over event from an avian reservoir host species^2^. The pathogen is now known to cause both upper and lower respiratory tract infections, including both bronchiolitis and pneumonia, with children being the most affected demographic^3–6^. It is estimated that 5% to 12% of hospitalizations due to viral respiratory infections of children are caused by HMPV^3, 7^. Furthermore, infections with HMPV can also be severe in the elderly and in patients with comorbidities^8^. Serology studies carried out in 2001 found that by the age of 5, virtually all children had been exposed to the virus^1^. Currently, there are no licensed vaccines or specific therapeutics for the treatment of HMPV infections. A better understanding of the molecular mechanisms of the HMPV life cycle are therefore needed.

HMPV shares many features with and is a close relative of respiratory syncytial virus (RSV), both being non-segmented negative sense RNA viruses (nsNSVs) belonging to the *Pneumoviridae* family^9^. HMPV possesses a 13.3 kb negative-sense RNA genome containing eight genes: 3’-N-P-M-F-M2-SH-G-L-5’^10^. The F, SH, and G proteins constitute surface glycoproteins^11–13^. The M and M2 genes encode the matrix protein M, and the M2-1 and M2-2 proteins via overlapping ORFs^14–16^. The L protein harbours the RNA-dependent RNA polymerase activity (RdRP) and is responsible for transcription of capped and polyadenylated viral mRNAs as well as replication of the genome^17^. The viral genome is encapsidated in a sheath of oligomerized copies of the nucleoprotein N, forming a ribonucleoprotein complex termed nucleocapsid^18, 19^. The nucleocapsid is the template for transcription and replication by L, which is also dependent on the obligate polymerase cofactor, the phosphoprotein P, together forming the active L/P holoenzyme^20^.

A hallmark of nsNSVs is the formation of specialized viro-induced cytoplasmic inclusions which concentrate viral RNA and proteins ^21^. During RSV and HMPV infections, these inclusion bodies (IBs) have been shown to harbour L/P and N and to be active sites of viral transcription and replication^22, 23^. *Pneumoviridae* IBs are spherical membraneless organelles. Recent studies provide evidence that IBs are biomolecular condensates that form through liquid-liquid phase separation (LLPS)^24–26^. HMPV IBs mature in the course of infection and grow via actin-dependent coalescence and fusion of replicative sites^23^. In a cellular context, the minimal elements for the morphogenesis of pneumoviral condensates are the P and N proteins, which together can form pseudo-IBs, even in the absence of any other viral proteins^25^. Of note, although the interaction between N and P has been demonstrated to be crucial for the formation of cellular IBs, HMPV P can phase-separate independently *in vitro*^27, 28^.

The nucleoprotein N is a promiscuous RNA-binding protein with globular N-terminal and C-terminal domains (NTD and CTD, respectively)^18, 29^. In oligomerized and RNA-bound N (N-RNA) the nucleic acid is threaded along an extended groove in between the NTDs and CTDs of laterally connected N protomers. Furthermore, N possesses short N-terminal and C-terminal extensions (“arms”) which contact neighbouring N protomers and facilitate oligomerization. The HMPV P protein is modularly disordered and forms a tetramer via a central coiled coil domain^30, 31^. Upstream and downstream of the coiled coil, P possesses extended regions which are mainly intrinsically disordered or present transient secondary structures in the absence of binding partners, but can conditionally fold upon interaction with other proteins. For instance, a recent cryo-EM reconstruction of the HMPV L/P complex showed contextual folding of regions of P lying downstream of the coiled coil through interaction with the polymerase^20^. However, in the same structure only 22% of the total residues of tetrameric P present in the construct were visible in the cryo-EM density, with the remainder being disordered. These substantial conformational dynamics make structural elucidation projects involving P challenging.

An important function of P is to bind to assembled N-RNA, forming an N-RNA/P complex. In both HMPV and RSV this occurs via an interaction of the disordered C-terminal region of P (P_CT_) with N^27, 28, 32^. The N-RNA/P interaction is thought to be important in multiple steps of the viral replicative cycle: 1) it tethers the L/P complex to the nucleocapsid while traversing the genome during transcription/replication, 2) it is required for the formation of IBs, 3) it is implicated in loading nucleocapsids into assembling progeny virions through an interaction with the viral matrix^33^. Even though these constitute crucial functions, the structural basis of this interaction in HMPV has thus far remained elusive.

In this study we report cryo-EM structures of recombinant HMPV N-RNA bound to P_CT_ (N-RNA/P), revealing the structural basis of the interaction surface. P binds into a groove at the periphery of the NTD of N, between a helix and an extended loop of a beta-hairpin of N (here denoted as N_βHL_). We observed that the density of N_βHL_ becomes more ordered upon P binding, indicative of a folding-upon-binding event. All-atom explicit solvent MD simulations of an N-RNA/P 5-mer indicated substantial conformational dynamics of the P_CT_, in line with the notion of a transient interaction. We further investigated pseudo-IB formation via a functional minigenome assay to quantify the effects of mutations within the beta-hairpin loop. Our experiments confirmed that N_βHL_ is important for binding of P and that perturbations of this interaction disrupt polymerase activity and IB formation. Taken together our data elucidate the structural mechanism of P attachment to N-RNA which retains substantial conformational flexibility in the bound state. Our observations are in line with a dynamic N-RNA/P interaction which allows rapid binding and unbinding during viral transcription/replication, yet is still sufficiently strong to maintain association through the multivalent character of the interaction partners.

## Results

### Cryo-EM reveals plasticity of N-RNA assemblies

*Pneumoviridae* nucleocapsids are helical assemblies of N bound to the viral RNA genome. Our strategy to elucidate the interaction between assembled N-RNA with the polymerase cofactor P was to employ a tractable system of recombinant N-RNA rings, described previously^29^. Viral N-RNA rings are present in both infected cells as well as in virions, however their function is currently unclear^34, 35^. We reasoned that recombinant N-RNA rings could be leveraged as a tool to obtain high resolution cryo-EM structures of a complex with P, circumventing confounding factors such as the inherent flexibility of helical N assemblies in *Pneumoviridae*^18^. As an initial step, we set out to determine the suitability of N-RNA rings for high resolution single particle cryo-EM. HMPV N-RNA rings were expressed and purified via size exclusion chromatography (Fig. 1A), followed by flash vitrification for cryo-EM. Raw EM images suggest a heterogenous mixture of multiple species. Template-free 2D-classification of particles revealed that the sample contains primarily rings with 10 protomers (Fig. 1B). However, 11-mer oligomers (black arrow, Fig. 1B), 12-mer oligomers (white arrow, Fig. 1B), and short coil-like segments, denoted as spiral particles, are also present in the preparation (red arrow, Fig. 1B).

**Fig. 1:**
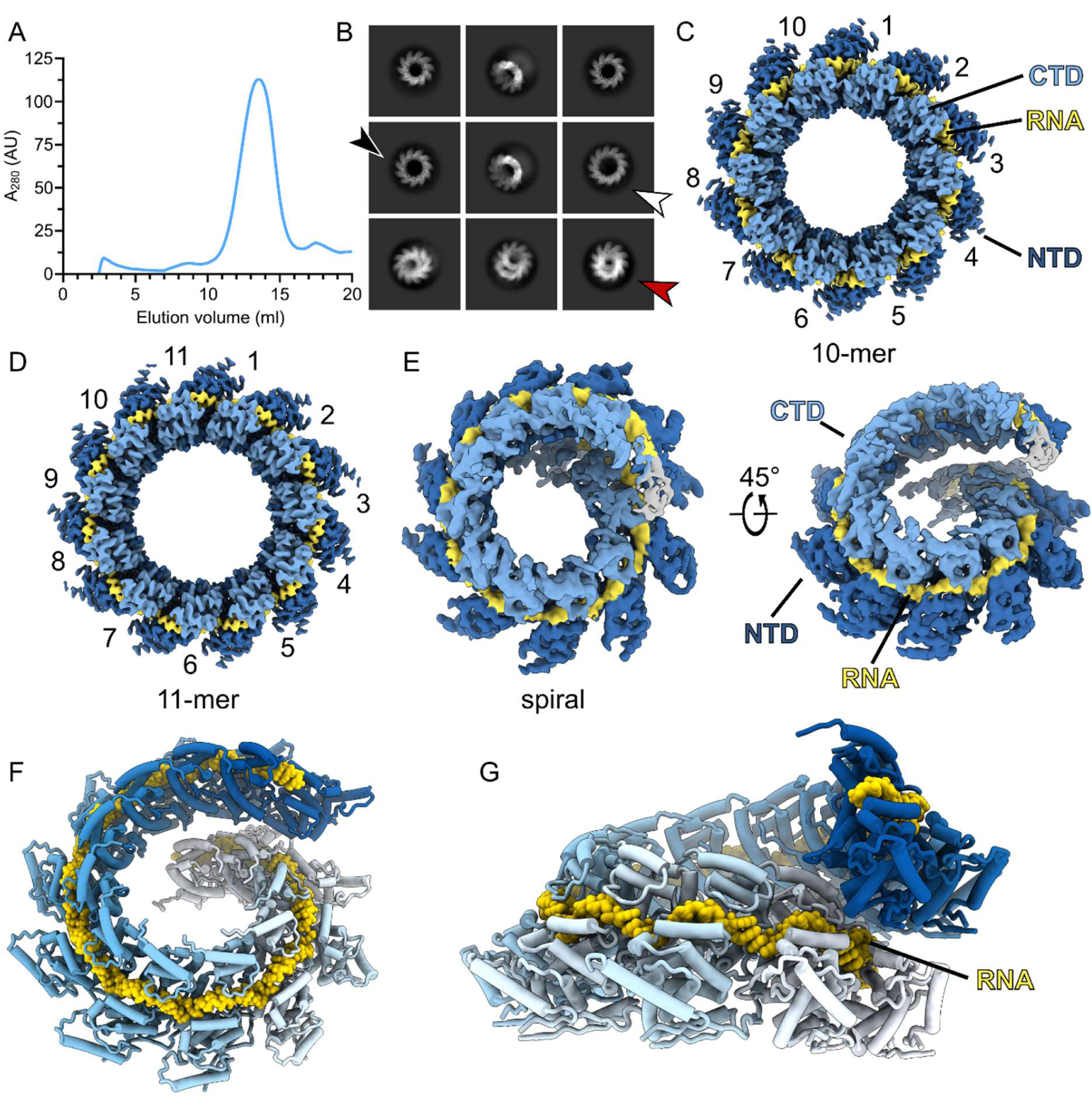
Cryo-EM of N-RNA from HMPV. (A) Size-exclusion chromatogram of recombinant HMPV N-RNA purified over a Superose 6 column. (B) Representative 2D-class averages from cryo-EM. Particles were found predominantly as 10-mer oligomers. However, 11-mers (black arrow), 12-mers (white arrow), and spiral particles (red arrow) were also observed. (C-E) Coulomb potential maps of single particle reconstructions of the indicated N-RNA oligomers. The bound RNA, and the C-terminal and N-terminal domains of N are labeled (CTD and NTD, respectively). The RNA is colored in yellow, the CTD in light blue, and the NTD in dark blue. (F) Tilted view and (G) side view of rigid body fitted N-RNA protomer models of the spiral assembly. N is shown in a cartoon representation, RNA is displayed as yellow spheres. The N protomers are colored sequentially from white to blue.

To determine if structures of suitable resolutions can be obtained from the heterogenous sample, we turned to single particle 3D reconstruction (Supplementary Table 1 and Supplementary Fig. 1A). We obtained a 3.1 Å resolution map of 10-mer rings of N-RNA (Fig. 1C and Supplementary Fig. 1B), a significant improvement from the previously reported crystallographic 10-mer at 4.2 Å^29^. We observed that the overall map quality was slightly better in the CTD region of N than in the NTD region, hinting at a degree of conformational heterogeneity in the NTD. we also determined a map of the 11-mer ring at 3.3 Å resolution (Fig. 1D). A structural alignment of the refined N protomer models solved by cryo-EM and crystallography shows that these are highly similar (Supplementary Fig. 2A, r.m.s.d. of 0.5 Å). The 11-mer of N-RNA possesses a diameter of ∼188 Å, slightly larger than the 10-mer at ∼180 Å (Supplementary Fig. 2B). The ring expansion by one protomer is accompanied by a slightly increased angle between neighboring N copies in the 11-mer (Supplementary Fig. 2C): 73.6° in the 11-mer vs 72.0° in the 10-mer, in respect to the center of the rings. In the case of the 12-mer the dataset did not contain a sufficient number of side-views to obtain a reliable map. Finally, 3D-reconstruction of the spiral particles yielded a coulomb potential map at an intermediate resolution of 4.7 Å (Fig. 1E).

The map density of the spiral particle shows N protomers forming a spiral staircase and stacking up upon each other, in a mode reminiscent of a helical assembly. The map quality is most reliable in the central protomers and becomes increasingly disordered towards the outer protomers. We were able to rigid body fit 12 protomers of N into the map (Fig. 1F). Thresholding of the map at a low level indicates the presence of additional N protomers, however this density was too diffuse to fit more copies of N, presumably due to heterogeneity of the sample or conformational heterogeneity. It is notable that in this arrangement the RNA strand is obscured by N protomers belonging to the next turn of the assembly (Fig. 1G), suggesting that a conformational change is necessary for the genome to become accessible to L/P. Taking together our cryo-EM reconstructions, we observe that the lateral interaction of RNA-bound N protomers can support a range of geometries, demonstrating the plasticity of N-RNA and supporting the notion of a highly flexible HMPV nucleocapsid.

### N protomers within assemblies are laterally hooked together via a loop insertion

While the previous analysis demonstrated that N has considerable flexibility in the types of N-RNA assemblies it can form, we wanted to also explore the dynamics of N protomers within an assembly. For this we turned to 3D Variability Analysis^36^, a methodology that can capture a family of related structures present in the experimental data. We decided to apply this to two neighboring N protomers of a 10-mer, following 5-fold symmetry expansion and local refinement. We observed that local refinement of an N-RNA dimer yielded higher quality maps (Fig. 2A) as evidenced by the improved anisotropic displacement parameters (ADPs) of the model (Supplementary Table 1), especially in the NTD region. Variability analysis of the dimer revealed a notable motion, especially of the NTD, given by a marked transverse tilting movement of N by ∼4°, in respect to the center of mass of the protomer (Supplementary Movie 1, Fig. 2B). The tilting motion occurs around a pivot point between N-terminal and C-terminal domains of N. As a consequence of the motion, the N-terminal domains globally move away from each other by ∼2 Å (maximum displacement up to ∼5 Å at the periphery of the NTD) at the endpoint vs. the starting point of the variability component.

**Fig. 2:**
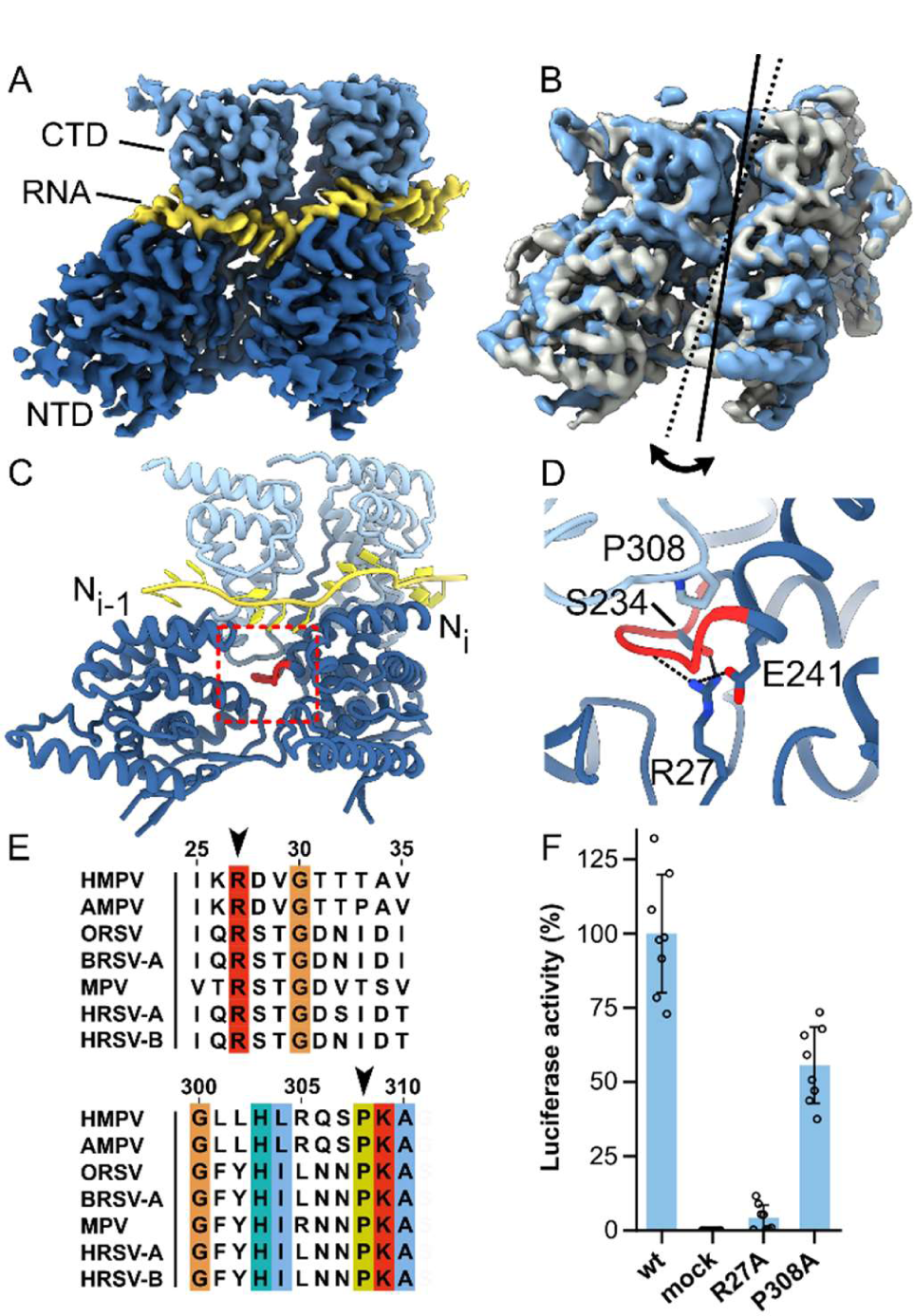
N protomers are hooked together via an inserted loop. (A) Representative cryo-EM density from local refinement. For clarity, only the map surrounding two N-RNA protomers is depicted. Domains of N and the bound RNA are labelled. (B) 3-D variability analysis (3DVA) was carried out with the N-RNA dimer. The density maps of starting point and end point of the first variability component are shown in grey and light blue, respectively. A transverse tilting motion could be observed within assembled N-RNA protomers (indicated with continuous and dotted axes). (C) Refined model of the N-RNA dimer. A loop (colored in red) of the Ni protomer is inserted into the neighboring Ni-1 protomer. (D) Close-up view of the region marked with a red square in panel C. Residues are shown in stick representation. The inserted loop is clamped from both sides via P308 and R27 from Ni-1. S234 and E241 further stabilize the interaction through H-bonding. (E) Multiple sequence alignment (MSA) of N sequences from *Pneumoviridae*. The positions of R27 and P308 are indicated with arrows. Abbreviations and UniProt accession codes: HMPV (human metapneumovirus, NCAP_HMPVC), AMPV (avian metapneumovirus, NCAP_AMPV1), ORSV (ovine respiratory syncytial virus, NCAP_ORSVW), BRSV-A (bovine RSV type A, NCAP_BRSVA), MPV (murine pneumonia virus, NCAP_MPV15), HRSV-A (human RSV type A, NCAP_HRSVA), HRSV-B (human RSV type B, NCAP_HRSVB). (F) HMPV polymerase activity in the presence of N mutants R27A and P308A measured by minigenome assay in BSRT7/5 cells. Data points represent the values of quadruplicates from two independent experiments. Error bars indicate the standard deviation.

The intra-assembly tilting of N begs the question of which N-N interactions at the lateral flanks remain constant and support side-to-side binding throughout the motion. While the importance of the N-terminal and C-terminal “arms” of N has been well established in this role^29^, we observed an additional mechanism of lateral N-N interaction. In all frames of the variability component, the loop ranging from residues 232-239 of the N_i_ protomer inserts into the neighboring N_i-1_ protomer (red loop in Fig. 2C). The loop of N is hooked into position from the top and bottom by residues P308 and R27 from the N_i-1_ protomer, with residues S243 and E241 providing additional stabilization through H-bonding (Fig. 2D). Multiple sequence alignment (MSA) of *Pneumoviridae* N proteins reveals that the proline and arginine residues at these positions are highly conserved (black arrows in Fig. 2E). To probe the importance of these residues for the replication/transcription of HMPV we employed a minigenome assay with a luciferase-based readout of viral polymerase activity^28^. Mutagenesis of P308 to alanine resulted in a notable decrease of polymerase activity, while the R27A mutation nearly abrogated polymerase activity (Fig. 2F), suggesting that the N-N interaction via the 232-239 loop is important for successful N-RNA assembly. Comparable expression levels of wild-type and mutant N proteins were observed in Western lots throughout the minigenome assays (Supplementary Fig. 3) In summary, our data points towards a critical lateral interaction between N protomers via insertion of the loop from residues 232-239, which stays consistent throughout the motion of assembled N protomers. Our data also suggest that residues P308 and R27 play a key role in this lateral interaction.

### The P_CT_ binds to the periphery of the NTD of N

Next, we set out to resolve the interface between the C-terminal region of P (P_CT_) and N-RNA. P is a flexible protein with extensive intrinsically disordered regions, and the N-RNA/P interaction has been suggested to be transient, complicating structural characterization^31, 32, 37^. Using the deep-learning based Metapredict server^38^, we generated disorder-score plots for both HMPV N and P. In agreement with previous work, N disorder scores are globally low, while P contains significantly disordered regions (Fig. 3A). The disorder prediction of the very C-terminal stretch of HMPV P lies in between ordered and disordered, suggestive of conditional disorder (black arrow in P disorder plot, Fig. 1A). Given the above considerations, our strategy was to shift the equilibrium to a P-bound state of N-RNA by adding a large molar excess of a P_CT_ peptide. We reasoned that the loss of contrast in the cryo-EM images stemming from unbound peptide background would not hinder us in high resolution 3D reconstruction due to the prominent and easy-to-align shape of N-RNA rings.

**Fig. 3:**
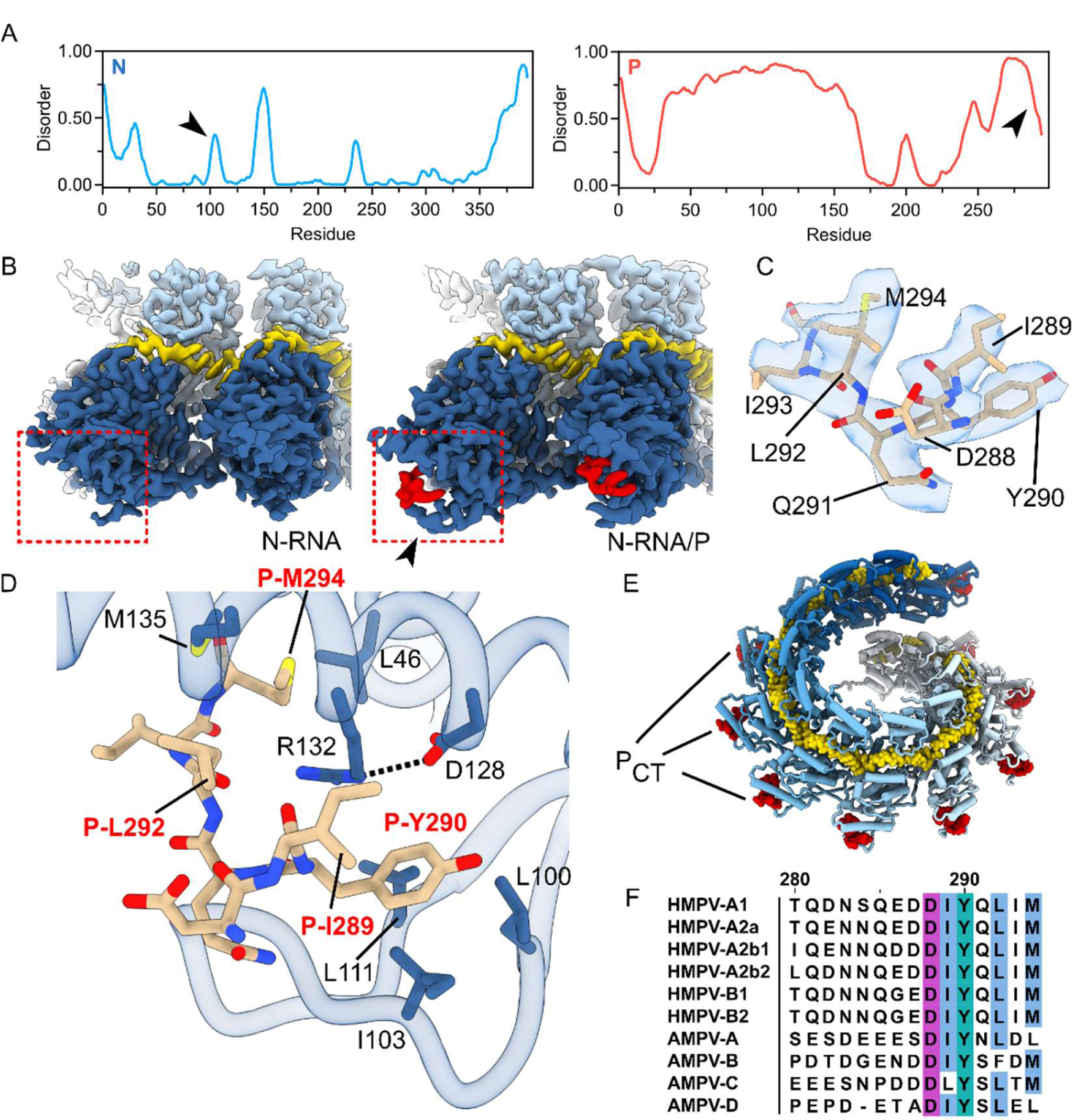
Cryo-EM structure of the N-RNA/P interface. (A) Disorder prediction plots of HMPV N (blue, left) and P (red, right) generated with Metapredict. Arrows, see accompanying text. (B) Comparison of locally refined cryo-EM maps before (left) and after (right) incubation with the PCT. The binding region is highlighted with a red square. The density of the PCT is colored in red. An increased ordering of density is observed in N residues 99-112 after addition of PCT (black arrow). (C) Density of the PCT with refined atomic model. The cryo-EM density is displayed as transparent surface, the model is shown in stick representation with labeled of P residues. (D) Close-up view of the PCT binding site. N residues are shown as blue sticks with black labels. P residues are shown as brown sticks with red labels. (E) Model of N-RNA/P protomers rigid body fitted into the 3D reconstruction of spiral particles incubated with PCT. PCT is labeled and depicted as red spheres. (F) MSA of P sequences from different genotypes of human and avian metapneumoviruses (HMPV and AMPV, respectively). GenBank accession codes: HMPV-A1 (AAK62967.1), HMPV-A2a (AGH27116.1), HMPV-A2b1 (AGJ74042.1), HMPV-A2b2 (QGV13025.1), HMPV-B1 (QTW05438.1), HMPV-B2 (QTW05474.1), AMPV-A (AAT68644.1), AMPV-B (AYO90670.1), AMPV-C (AHA36959.1), AMPV-D (CDN30035.1).

We incubated N-RNA preparations with a ∼100-fold molar excess of synthetic peptide covering the 9 last residues of P_CT_ (N-EDDIYQLIM-C). Cryo-EM analysis of the N-RNA/P complex revealed the presence of the same classes of assemblies as was the case with the apo sample. We carried out 3D reconstruction of 10-mers, 11-mers, and spiral particles of the N-RNA with P_CT_ (Supplementary Table 1, Supplementary Fig. 4). In the reconstructions, additional density was visible at the periphery of the NTD of N. To facilitate easier model building and to obtain improved maps, we carried out symmetry expansion and local refinement of a dimer of N-RNA/P as before with N-RNA. The final resolution was at 2.9Å and comparison of the N-RNA and N-RNA/P dimer maps (Fig. 3B) revealed density of sufficient quality to allow refinement of a model of the bound P_CT_ (Fig. 3C).

The cryo-EM map resolved the residues 288 to 294 of the P_CT_ bound to N-RNA. P_CT_ is wedged in between a helix of N (N residues 121-141) on one side and an extended loop of a beta-hairpin of N (N_βHL_, N residues 99-112) on the other, burying a total of area ∼1100 Å^2^ upon complex formation. We observed a noticeable increase of ordered density of N_βHL_ upon P-binding, compared to the apo form (black arrow, Fig. 3B). Of note, N_βHL_ is also predicted to be slightly disordered in our Metapredict plots (black arrow in N disorder plot, Fig. 1A). We observe that the P peptide wraps around a central arginine (R132) positioned in the middle of the helix of N (Fig. 3D). P hooks onto this arginine by packing the hydrophobic surfaces of P_CT_ side-chains P-M294, P-L292, P-I289, and P-Y290 against the aliphatic chain of R132, thereby completely surrounding it (Fig. 3D). R132 is additionally stabilized and kept in position through a polar contact with D128. The C-terminal methionine of P (P-M294) inserts into a hydrophobic cavity on N formed by L46 and M135. Moreover, the face of the aromatic ring of P-Y290 packs against three hydrophobic residues located within N_βHL_: L100, I103, and L111 (Fig.3D). Lastly, viewed in the context of an N-RNA assembly, the P_CT_ binding site is located such that P attaches to a region of the nucleocapsid that protrudes at a far radius from the helical axis (Fig. 3E). This may facilitate accessibility to P-binding, even in light of the conformational heterogeneity of the nucleocapsid.

Past studies have utilized functional and biochemical assays to investigate the HMPV N-RNA/P interaction^27, 28^. Our structural observation of the importance of P residues P-M294, P-L292, P-Y290, and P-I289 for the interaction with N-RNA is fully in line with the previous studies, which determined that single substitutions of these residues (compare with Fig. 3D) to alanine completely disrupted the interaction. The same observation could be made with substitutions of N residues R132 and D128, confirming our structure and indicating that D128 is important to orient R132 in a manner that facilitates P-binding^28^. In addition, a MSA incorporating P_CT_ sequences from metapneumovirus members shows the conservation of hydrophobic residues in positions 294, 292, 290, and 289 (HMPV numbering), highlighting their importance in clamping onto N (Fig. 3F).

To investigate the dynamics of P_CT_ bound to N-RNA we carried out all-atom explicit-solvent MD simulations of the triple complex. To circumvent artifacts stemming from the simulation of individual protomers lacking neighbors, we simulated 5-mers of assembled N-RNA/P. The overall conformation of the N-RNA protomers was consistent throughout the simulations (Fig. 4A). The RNA remained tightly bound, with low root-mean-square-deviations (RMSDs) and low fluctuations in the number of RNA-protein hydrogen bonds over the simulation times (Fig. 4B). In contrast, substantially higher RMSD values were observed for many of the copies of P-peptide bound to N (Fig. 4C) and the peptide displayed large conformational flexibility (black arrow in Fig. 4A and Fig. 4D). Notably, in one case a peptide completely dissociated from its starting N protomer (black arrow, Fig. 4C), and re-bound at a neighboring copy of N, displacing an already present P_CT_ (Fig. 4E and Supplementary Movie 2). These data indicate that the interaction between P_CT_ and N-RNA is flexible and suggest that P has the ability to dynamically unbind and rebind on simulated timescales that are notably rapid (i.e. tens to hundreds of nanoseconds).

**Fig. 4:**
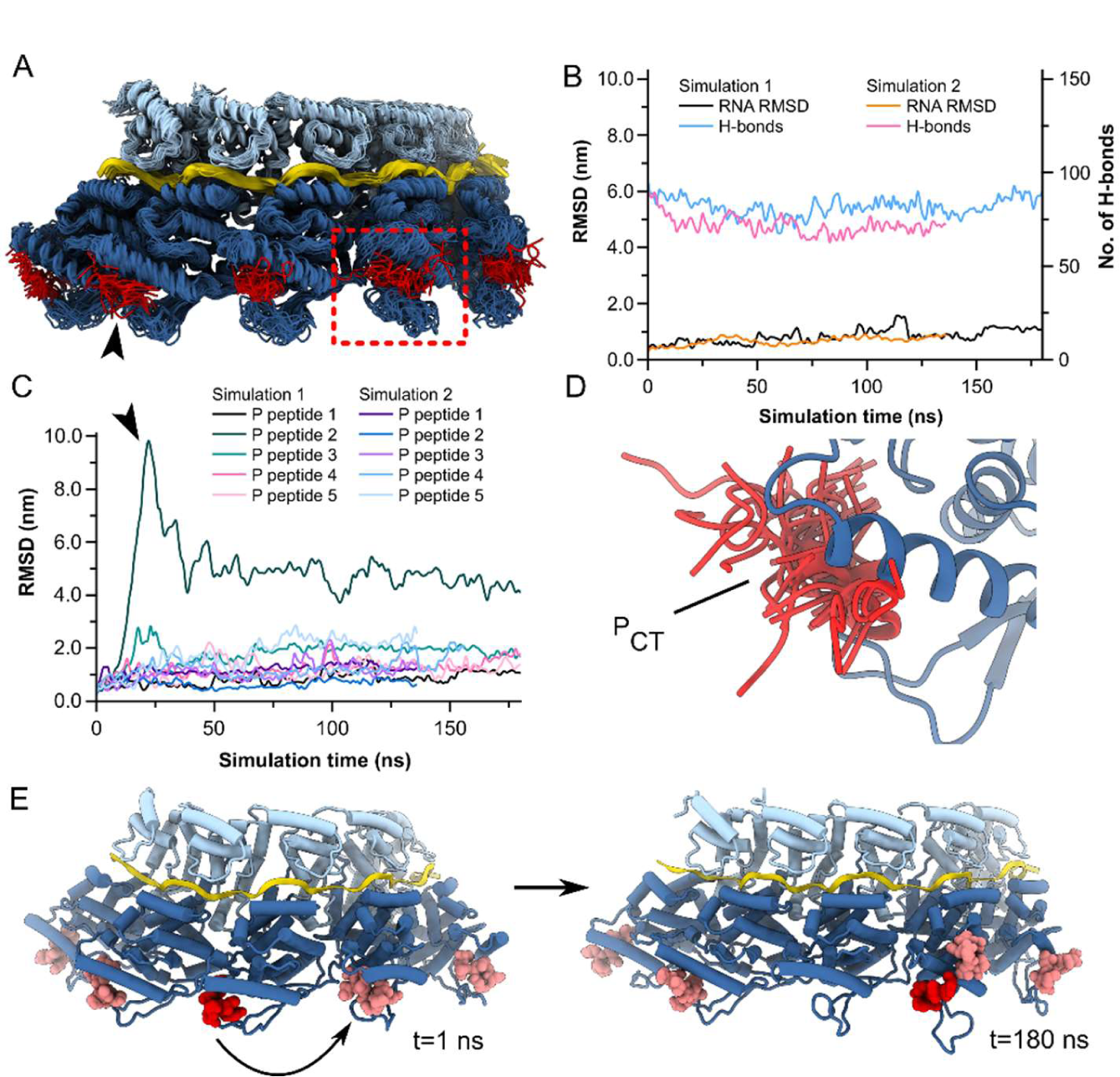
MD simulations of N-RNA/P. (A) Overlayed structural ensemble of a simulated trajectory of a N-RNA/P 5-mer. Snapshots were taken every 5 ns. PCT was observed to be highly dynamic, even in the bound state (black arrow). (B) Time evolution of hydrogen bonds and RMSD of bound RNA molecules over the simulation trajectories. Data are shown for duplicate simulations. (C) Time evolution of RMSDs of PCT peptide over the simulation trajectories. High RMSD values were observed for multiple peptides. In one case a peptide unbound and rebound at a neighboring N-RNA protomer (trajectory indicated by black arrow). (D) Ensemble of PCT conformations highlighting its flexibility. For clarity only one ensemble member is shown for N. (E) Snapshots of a simulated trajectory at t = 1ns (left) and t = 180 ns (right). PCT peptides are colored in shades of red. In the evolution of one trajectory a copy of the PCT (colored in dark red) unbinds, and rebinds at a neighboring N protomer, displacing an already present PCT (indicated with curved arrow).

### Interaction between N_βHL_ and P_CT_ is important for polymerase activity and IB formation

Next we investigated the functional role of the interaction between P_CT_ and the flexible N_βHL_ within cells using minigenome assays. We focused our mutations on the hydrophobic residues L100, I103 and L111 of N, which pack against Y290 of the P_CT_ (see Fig. 3C). Whereas mutations of residue I103 poorly impacted the polymerase activity, mutations of residues L100 and L111 resulted in a noticeable reduction of polymerase activity in the minigenome assay (Fig. 5A). We confirmed by Western blot that this was not due to large variations in expression levels of mutant N proteins (Supplementary Fig. 3) The strongest effect for a single substitution was observed for the leucine at position 111. Exchange to an alanine (L111A) resulted in a polymerase activity decrease to below 50% while an introduction of a negative charge (L111E) led to a decrease to ∼ 10% activity, compared to wild-type. A triple substitution mutant of L100, I103, and L111 to alanine reduced the activity to around a third (∼30%), while a corresponding triple mutant with substitutions to glutamic acid in these positions abrogated polymerase activity (Fig. 5A).

**Fig. 5:**
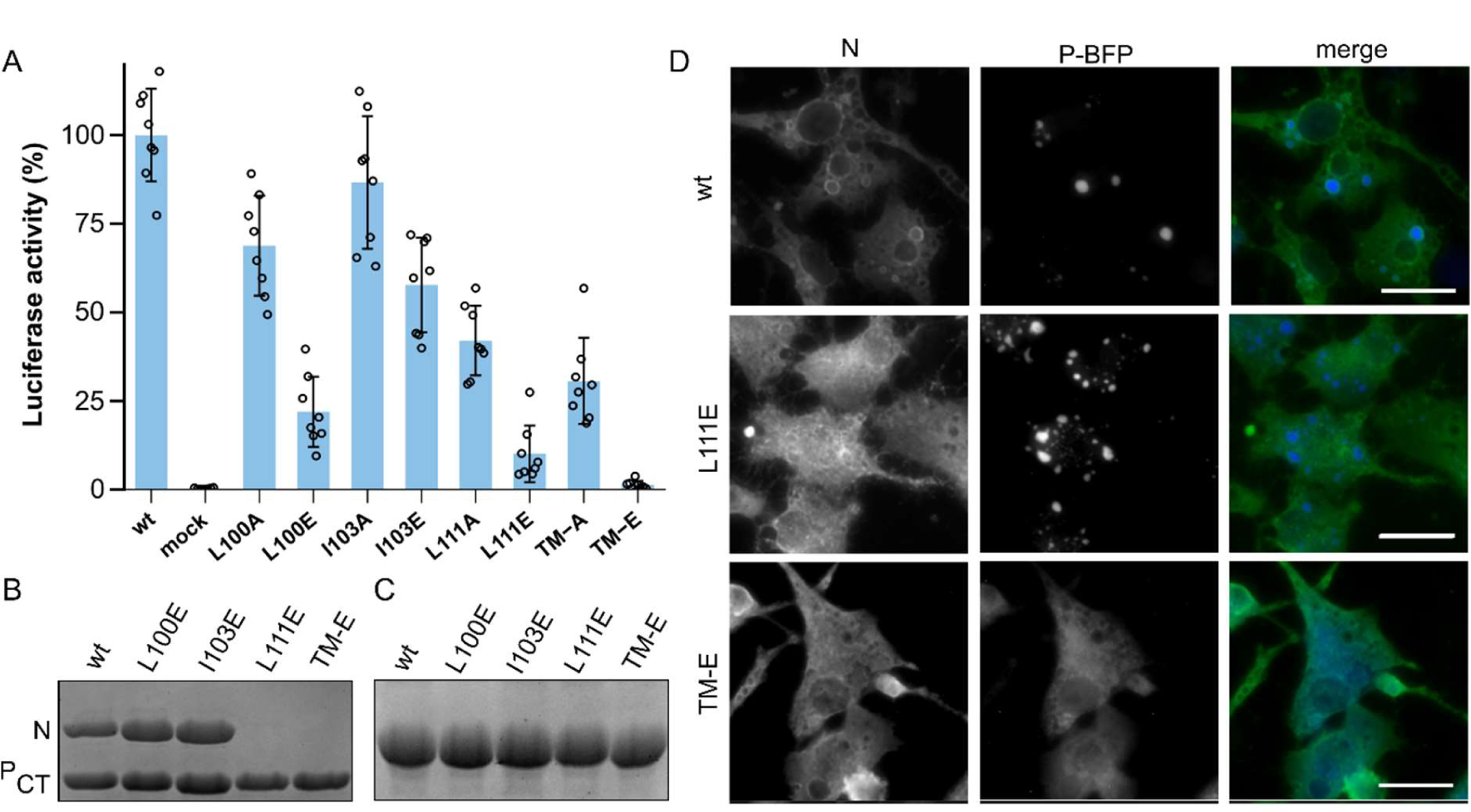
Functional role of the NβHL in PCT recognition. (A) HMPV polymerase activity in the presence of the indicated N mutants, assessed by minigenome assay in BSRT7/5 cells. TM-A denotes the triple substitution mutant L100A, I103A, L111A. TM-E denotes the triple substitution mutant L100E, I103E, L111E. Data points represent the values of quadruplicates from two independent experiments. Error bars indicate the standard deviation. (B) GST-PCT and 6xHis-tagged N proteins (wt and mutants) were co-expressed and co-purified using a GST-tag. A SDS-PAGE analysis of the products of purification is shown, with the position of N and GST-PCT indicated. (C) SDS-PAGE analysis of N mutants purified using 6xHis-tag, demonstrating that these are soluble. (D) Fluorescence microscopy of BSRT7/5 cells transfected with plasmids encoding BFP-tagged HMPV P and HMPV N (wt and mutant). Cells were fixed 24h post-transfection. N protein was revwealed by labelling with rabbit polyclonal anti-N antibody and nuclei were stained with Hoechst 33342. Scale bars, 20 µm.

Next, we explored the effect of the most impactful substitutions on the ability of N and P to physically interact or to form inclusion bodies (IBs). We first assessed the ability of a GST fused to P_CT_ to pull down N (wt or mutant) upon co-expression in bacteria(Fig. 5B, uncropped gels in Supplementary Fig. 5). While L100E and I103E retained the association with N, constructs possessing the L111E substitution were not able to copurify N. We also assessed that the loss of interaction was not due to a defect of N expression or solubility, by purifying the individual N proteins expressed in bacteria (Fig. 5C).

The expression of P and N is sufficient for the formation of pseudo-IBs in cells^24, 27^. To test the effect of N_βHL_ mutants on IB formation, we co-transfected BSRT7/5 cells with plasmids encoding N (wt and mutants) and BFP-tagged P and imaged the cells by fluorescence microscopy. The P-BFP fluorescence allowed us to observe pseudo-IBs, while labelling with anti-N antibodies allows only the observation of inclusion outlines, likely due to inaccessibility of the interior. Compared to the wt condition, pseudo-IBs formed in the presence of L111E mutant of N were more numerous and smaller, suggesting a defect of nucleation. In line with the pull-down and minigenome assays, the N triple mutant TM-E resulted in a total loss of IB formation, with the P protein being distributed throughout the cytoplasm. Together, these results confirm that N_βHL_ is in contact with P, and that perturbation of this contact leads to a loss of polymerase activity and affects LLPS induced by the interaction of N and P proteins.

## Discussion

The viral RNA genome of all nsNSVs is constitutively encapsidated by the nucleoprotein N, forming the nucleocapsid. Packaging within this ribonucleoprotein complex is thought to protect the genome from degradation by cellular RNases but also from the recognition by sensors of the cellular immune system such as RIG-I. However, a consequence of the encapsidation is that it is not naked RNA but rather the N-RNA complex which serves as a template for the viral polymerase L. The P protein is an essential cofactor of L, which interacts with both L and N-RNA, functioning as a flexible tether. The recent cryo-EM structures of *Pneumoviridae* L/P complexes revealed that the central oligomeric domain of the P tetramer and the four C-terminal domains of tetrameric P interact with L^20, 39, 40^. These structures also showed that each of the C-terminal domains of P adopt specific and different conformations upon interaction with L, highlighting the structural versatility of P which is required for the proper functioning of L/P. Of note, in these structures the last C-terminal residues of P, identified as essential for the interaction with the nucleocapsid^27, 28^, were not resolved. The structures imply that one tetramer of P can interact with both L and the nucleocapsid simultaneously. Residues of HMPV P and N involved in the P-nucleocapsid interaction have been identified^27, 28^. Previous crystallographical studies of this N-P interaction in the related pneumovirus RSV utilized a truncated N protein and revealed density for only the last two residues of P^32^. The results of this study suggested that the C-terminus of P may be flexible upon binding N, which agrees with the weak strength of this interaction (in the micromolar range)^32^. Previous attempts to elucidate how L/P attaches to N-RNA thus remain limited, and the impact of P binding on full length N is still unknown.

Here, using cryo-EM and full-length N, we were able to determine the structures of assembled and RNA-bound HMPV N in different oligomeric states, including a spiral particle reminiscent of a helical turn, and with improved resolution compared to a previous crystal structure^29^. Analysis of our structural data revealed heterogeneity in the relative geometries of assembled N protomers, supporting the notion of a highly flexible nucleocapsid. The observed dynamics of N protomer orientations, even in the assembled state, may constitute a prerequisite for conformational changes facilitating access to the RNA genome during transcription by L/P. Furthermore, we identified a key N-N interaction involving the loop ranging from residues 232-239 of the N_i_ protomer which inserts into the neighboring N_i-1_ protomer. We found that residue R27 of the loop was sensitive to mutation and a single exchange to alanine had a major impact on polymerase activity. However, further structural characterization of N-RNA oligomers of the R27A mutant are necessary to determine the precise role of this interaction for nucleocapsids.

We were successful in obtaining the first structural data on the HMPV P_CT_-N-RNA interaction. Utilizing an approach that made use of a large molar excess of P_CT_ vs. N, computational symmetry expansion, and local refinement we were able to resolve P_CT_ bound to the NTD of assembled and RNA-bound N. Our structure reveals how P binds by hooking itself around a centrally positioned arginine of the NTD, thereby latching on to N-RNA. The P_CT_ interacts with the partially disordered N_βHL_ region of the NTD, which becomes more ordered upon binding. Furthermore, in the apo-state the P_CT_ itself has been characterized as having high disorder propensity^30, 31^. Such folding upon binding events of intrinsically disordered regions are thought to facilitate highly specific interactions, yet possess low affinities and rapid dissociation kinetics^41^. We acknowledge that our experimental conditions involving ∼100-fold molar excess of P_CT_ would not be the case *in vivo,* but likely assisted in shifting the equilibrium to a bound state, enabling resolving the site.

To explore the dynamics of the P_CT_ interaction with N we turned to all-atom MD simulations of 5 protomers of N-RNA/P. The simulations revealed substantial conformational flexibility of P_CT_ within the N-RNA/P complex. In one instance we observed the dissociation and rebinding of one of the simulated peptides. This is in line with the disordered character of P, but also with previous reports in RSV characterizing the interaction as giving rise to a fuzzy complex^37, 42^.

Taking together the properties of the N-RNA/P interaction, we suggest that its evolution is constrained by a need to balance sufficient tethering of L to the nucleocapsid with being able to unbind/rebind with low energy barriers during transcription. RNA-polymerases exert pulling forces in the tens of Piconewtons range^43^ and in the case of L/P this needs to be sufficient to dislodge tetrameric P from any given position on the nucleocapsid while it is being traversed by L/P. In the hypothetical case of a P_CT_ binding too tightly to N-RNA, the pulling force might not be able to overcome this tethering, leading to a stalled complex. At the same time, the interaction needs to be strong enough to recruit L/P for transcription. Several decades ago a “cartwheeling” model was proposed for the Sendai paramyxovirus^44^, suggesting that “P is envisaged to “walk” on the template via the simultaneous breaking and reforming of subunit-template contacts”. Our data fully support the applicability of this classic model for the *Pneumoviridae* L/P complex, in which it scuttles along the nucleocapsid template during processive RNA synthesis using the C-terminal domains of tetrameric P as legs.

Our study provides a structural blueprint for a potentially druggable interface in HMPV. Previous attempts have been made to target the N/RNA-P interaction in RSV, resulting in the compound M76, which insert into the P_CT_ binding site on RSV N^32^. Unfortunately, antiviral inhibition through M76 was limited. More recently, the 1,4-benzodiazepine-derived compound EDP-938 shows promising antiviral activity against RSV-A and RSV-B, a relatively high barrier against resistance, efficacy in non-human primates, and was superior to placebo in a randomized, double-blind challenge trial^45, 46^. As of 2023, EDP-938 is still undergoing phase II clinical trials. EDP-938 has been shown to inhibit a post-entry step of the viral replicative cycle and reduce the accumulation of viral RNA early in the infection^45^. Based on mapping of resistance mutations, it was concluded that the compound targets the RSV N protein, although the mechanism of inhibition has not yet been finally settled. Interestingly, the by-far strongest resistance mutant to EDP-938 (M109K, responsible for a 67-fold worsening of the EC_50_) is located in a region corresponding to the N_βHL_ of HMPV (red spheres in Fig. 6A). Comparison with our N-RNA/P complex (Fig. 6A,B), with its extended interaction surfaces, suggest that M109 may be involved in transiently clamping down on the P_CT_ in RSV, in analogy to HMPV. This is also in-line with our functional data which show that N_βHL_ mutations modulate the interaction with the peptide. These observations strengthen the hypothesis that the N-RNA/P interaction may be the target of EDP-938. However, the matter is complicated by the fact that EDP-938 had no effect on transcription in a minigenome assay^45^. Whether this may be due to experimental constraints or because there are additional unknown factors in the mechanism of the compound will need to be explored in future work.

**Fig. 6:**
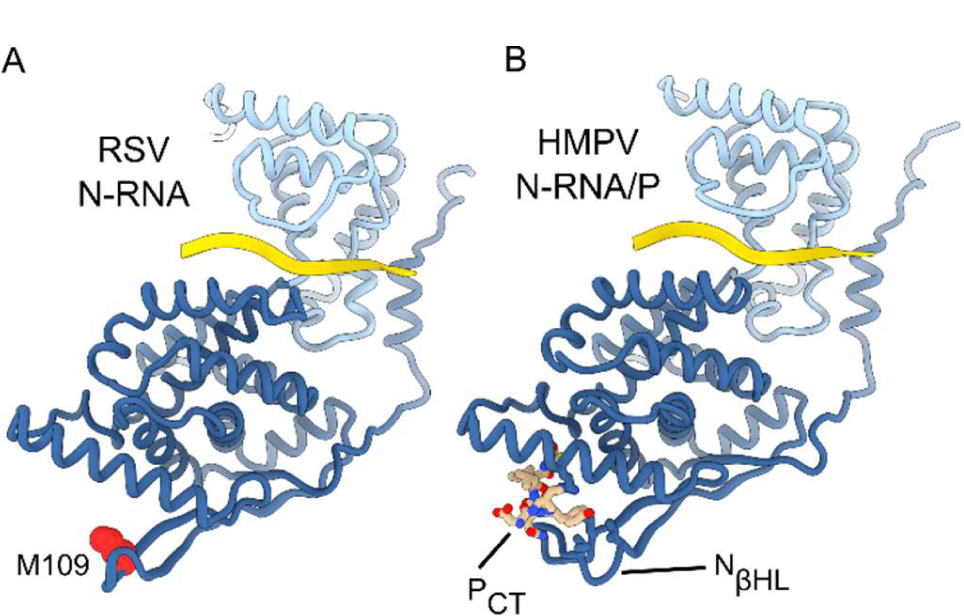
Comparative analysis of N-M109 resistance mutation against EDP-938 in RSV. (A) Structure of a HMPV N-RNA/P protomer. The PCT is indicated and depicted as sticks. (B) Structure of a RSV N-RNA protomer (PDB ID: 2wj8). The residue M109 is indicated and depicted as red spheres.

## Methods

### Plasmids for pulldowns and minigenome assays

The plasmids pET-N and pGEX-P_CT_ encoding for N with a C-terminal 6xHis tag and for the C-terminal residues 200-294 of P fused to GST respectively, with N and P sequences derived from HMPV CAN97-83 strain, were described previously^28^. The HMPV minigenome plasmid containing Gaussia/Firefly luciferases and the plasmids pP, pN, pL, and pM2-1 corresponding to the sequences of NL99-1 HMPV strain cloned into pCite vector, as well as pP-BFP vector were also previously described^28^. Point mutations were introduced in the N sequence by site-directed mutagenesis using the Q5 site-directed mutagenesis Kit (New England Biolabs). Sequence analysis was carried out to check the integrity of all constructs. All the oligonucleotides sequences are available on request.

### Antibodies

Antisera used in this study included polyclonal rabbit antisera raised against recombinant HMPV N expressed in bacteria. A mouse monoclonal anti-β-tubulin (Sigma) and secondary antibodies directed against mouse and rabbit Ig G coupled to HRP (P.A.R.I.S) were also used for immunoblotting. Secondary antibody directed against rabbit Ig G coupled to Alexafluor-488 (Invitrogen) was used for immunofluorescence experiments.

### Cell culture and transfections

BHK-21 cells (clone BSRT7/5) constitutively expressing the T7 RNA polymerase^47^ were grown in Dulbeco Modified Essential Medium (Lonza) supplemented with 10% fetal calf serum (FCS), 2 mM glutamine, and antibiotics. The cells were grown at 37°C in 5% CO_2_, and transfected using Lipofectamine 2000 (Invitrogen) as described by the manufacturer.

### Minigenome replication assay

BSRT7/5 cells at 90% confluence in 96-well dishes were transfected using Lipofectamine 2000 (Invitrogen) with a plasmid mixture containing 62.5 ng of pGaussia/Firefly minigenome, 62.5 ng of pN, 62.5 ng of pP (WT and mutants), 31.25 ng of pL, 31.25 ng of pM2-1, as well as 15.6 ng of pSV-β-Gal (Promega) to normalize transfection efficiencies. Cells were harvested at 24 h post-transfection and lysed in 100 µl of Firefly lysis buffer (30 mM Tris [pH 7.9], 10 mM MgCl2, 1 mM dithiothreitol [DTT], 1% [vol/vol] Triton X-100, and 15% [vol/vol] glycerol). Then 50 µL of lysates were mixed to 50 µl D-luciferine (Luciferase assay system, Promega) and the Firefly luciferase activity was determined for each cell lysate with an Infinite 200 Pro (Tecan, Männedorf, Switzerland), and normalized based on β-galactosidase (β-Gal) expression. Transfections were done in quadruplicate, and each independent transfection experiment was performed up to four times. For proteins expression analysis, cells were lysed in Laemmli buffer, boiled, and analyzed by Western blotting (WB) according to standard protocols.

### Fluorescence microscopy

Immunofluorescence microscopy was performed with cells grown on coverslips and previously transfected with pP-BFP and pN (WT or mutants), at a ratio 1:1. At 24 h post transfection, cells were fixed with 4% paraformaldehyde (PFA) for 20 min, made permeable and blocked for 30 min with PBS containing 0.1% Triton X-100 and 3% bovine serum albumin (BSA). Cells were then successively incubated for 1 h at room temperature with primary and secondary antibody mixtures diluted in PBS containing 3% BSA and 0.05% Tween20. Coverslips were mounted with ProLong Gold antifade reagent (Invitrogen) and observed with an inverted fluorescence microscope (Zeiss Axiovision). Images were processed with ZEN software (Zeiss) and ImageJ software.

### Expression and purification of recombinant proteins

*E. coli* BL21 bacteria (DE3) (Novagen) transformed with pET-N (WT or mutants) alone or with pGEX-P_CT_ were grown at 37°C for 8 hours in 100 ml of Luria Bertani (LB) medium containing 100µg/ml ampicillin or ampicillin (100 µg/ml) and kanamycin (50 µg/ml), respectively. The same volume of LB was then added and protein expression was induced by adding 80 µg/ml isopropyl-ß-D-thio-galactoside (IPTG) to the medium. The bacteria were incubated for 15 hours at 28°C and then harvested by centrifugation. For purification using the GST-tag, bacteria were re-suspended in lysis buffer (50 mM Tris-HCl pH 7.8, 60 mM NaCl, 1 mM EDTA, 2 mM DTT, 0.2% Triton X-100, 1 mg/ml lysozyme) supplemented with complete protease inhibitor cocktail (Roche), incubated for 1 hour on ice, sonicated, and centrifuged at 4°C for 30 min at 10,000g. Glutathione-Sepharose 4B beads (GE Healthcare) were added to clarified supernatants and incubated at 4°C for 3 hours. Beads were then washed with 10 volumes of lysis buffer, then three times with PBS 1X, then stored at 4°C in an equal volume of PBS.

For purification of N-6xHis fusion protein purification, bacterial pellets were re-suspended in lysis buffer (20 mM Tris-HCl pH8, 500 mM NaCl, 0.1% TritonX-100, 10 mM imidazole, 1 mg/ml lysozyme) supplemented with complete protease inhibitor cocktail (Roche). After sonication and centrifugation, lysates were incubated 30 min with chelating Sepharose Fast Flow beads charged with Ni^2+^ (GE Healthcare). Beads were then successively washed in the washing buffer (20 mM Tris-HCl, pH 8, 500 mM NaCl) containing increasing concentration of imidazole (25, 50, and 100 mM), and proteins were eluted in the same buffer with 500 mM imidazole. For SDS-PAGE analysis, samples were prepared in Laemmli buffer, denatured 5 min at 95 °C, separated on 12% polyacrylamide gel colored by Coomassie brilliant blue.

### Expression and purification of HMPV N-RNA for structural analysis

The full-length N protein gene from human metapneumovirus (strain NL1-00, A1, GenBank: AAK62966.1) was cloned into the pOPINE bacterial expression vector, also coding for a C-terminal His-tag, using the In-Fusion system (Takara Clontech, Mountain View, CA) following manufacturer’s protocols^48^. The vector DNA sequence was verified by means of Sanger sequencing. Rosetta2 *E. coli* cells containing the plasmid were grown at 37°C in terrific broth containing appropriate selection antibiotics. N protein expression was induced once an OD_600_ of 0.8 was reached, by adding isopropyl β-D-1-thiogalactopyranoside to a final concentration of 1 mM. The temperature was then lowered to 18°C and cells were harvested after overnight growth (18°C, 20 min, 4000 x g). Cell pellets were resuspended in 40 mL of wash buffer (25 mM Tris-HCl, pH 8.0, 1 M NaCl) per litre of bacterial culture, and lysed by means of sonication. The resultant lysate was clarified (4°C, 45 min, 50000 x g) and the supernatant was filtered and loaded on a column containing Ni^2_^NTA (nitrilotriacetic) agarose (Qiagen, Netherlands). The column was equilibrated with wash buffer and the bound protein washed. The protein was eluted in elution buffer (25 mM Tris-HCl, pH 8.0, 1 M NaCl, 400 mM Imidazole), and the resultant eluate was further purified by size exclusion chromatography, using a Superose6 10/300 column (GE Healthcare, United Kingdom), equilibrated in SEC buffer (25 mM Tris-HCl, pH 8.0, 1 M NaCl). The protein was then buffer exchanged into 25 mM Tris-HCl, pH 8.0, 250 mM NaCl using a PD10 column (GE Healthcare).

### Preparation of N-RNA/P complexes

Synthetic peptides of P covering the sequence N-EDDIYQLIM-C were obtained from GenScript. The peptide was dissolved in deionised water dosed with a drop of 5 M NH_4_OH to improve solubility. N-RNA preparations were mixed with the resulting peptide solution in a ∼100-fold molar excess of P peptide to N. The N-RNA and PCT mixture was incubated at 4°C over night.

### Cryo-EM data collection and processing

Cryo-EM experiments were carried out at the Oxford Particle Imaging Centre (OPIC). 3 µL of N-RNA (1.0 mg/ml) or N-RNA/P (0.5 mg/ml) sample were placed onto glow-discharged Quantifoil R2/1 Cu300 holey carbon grids (1 µm spacing, 2 µm holes, copper mesh) (Quantifoil, Germany), blotted for 3.5 s before flash-freezing in liquid ethane using a Vitrobot mark IV (FEI) at 95 – 100 % humidity. Cryo-EM data were collected on a 300 kV Titan Krios microscope (Thermo Fisher Scientific) fitted with a K2 Summit (Gatan) direct electron detector and GIF Quantum energy filter. Data collection parameters are listed in Supplementary Table 1 for both datasets. Motion correction of cryo-EM movies and contrast transfer function (CTF) parameters were estimated ‘on-the-fly’ using cryoSPARC live^49^.

All single particle reconstructions were carried out in cryoSPARC. Particle picking was carried out using a combination of the cryoSPARC blob picker and template picker. 2D classification was utilized to separate particle subsets of different oligomeric states and remove bad particles. Particles were further cleaned through heterogeneous refinement. The final maps were obtained from the non-uniform refinement job type. Symmetry was imposed only when it was clearly visible in template-free 2D class averages, as was the case for the 10-mer and 11-mer classes (C10 and C11 symmetry imposed, respectively). No symmetry was imposed for the refinement of spiral particles. To improve the density of N-RNA protomers we carried out 5-fold symmetry expansion of the 10-mer particles, followed by local refinement of a N-RNA dimer. We initially attempted an approach with 10-fold symmetry expansion and local refinement of a N-RNA monomer, however this yielded worse maps, presumably because a single monomer is not of sufficient size for local refinement (less than 50 kDa). Total particle numbers and numbers of particles used for the reconstruction of every map can be seen in Supplementary Figs. 1 and 4 for N-RNA and N-RNA/P, respectively, and in Supplementary Table 1.

### Model building and refinement

The model of HMPV N obtained from X-ray crystallography was used as a starting point for further refinement (PDBID: 5fvc). Iterative cycles of manual refinement and building in COOT^50^ and real-space refinement in PHENIX^51^ were used to optimize geometry and fit to the density. PHENIX validation tools were used to monitor geometry and density fit parameters. For maps with imposed symmetry (10-mer and 11-mer maps) we refined a single protomer copy and generated symmetry mates for figure preparation. The N-RNA models (from maps without P) encompassed residues 3 to 365 of the N protein, with a gap from residues 100 to 111 where the density was not sufficient for building a model. Moreover, we observed only very incomplete and diffuse density after residue 365 (the C-arm) and thus refrained from continuing to build the model after this point. For the models of the spiral particles we limited ourselves to rigid body fitting of protomers (from the high resolution refinements), due to the low resolution of the spiral maps. For the N-RNA/P complex, the local refined map showed improved density in the region of residues 100-111 and thus allowed building a continuous model up to residue 365. The model also included P residues 288 to 294. The N-RNA/P model from the locally refined dimer maps was then used as a basis for the model refinement of N-RNA/P 10-mers and 11-mers, as well as for rigid body fitting into the N-RNA/P spiral map. Model refinement statistics are given in Supplementary Table 1.

### Molecular dynamics simulations

Classical MD simulations were used to study the dynamics of the N-RNA and N-RNA/P_CT_ complexes. A 5-mer of N-RNA complex, with or without the 5 bound P_CT_ peptides, was extracted from the N-RNA/P cryo-EM structure. The MD systems were set up using the solution builder tool from CHARMM-GUI input generator ^52^. Briefly, each system was solvated in a rectangular periodic box which size was determined by the biomolecular extent, resulting in the addition of approximately 200k TIP3P water molecules. The systems were neutralized by adding 150mM NaCl. The two systems were then energy minimized, equilibrated in NVT ensemble and simulated for ∼180ns in 2 independent trajectories in GROMACS2021^53^ by making use of the CHARMM-GUI provided scripts^54^. The CHARMM36m force field was used for all simulations^55^. The trajectories were analyzed using GROMACS tools to extract RMSDs and number of hydrogen bonds as a function of simulation time.

### Structure analysis and figure preparation

Structures were inspected and analysed using tools within UCSF Chimera and ChimeraX^56, 57^. Structure figures and movies were prepared in ChimeraX. Multiple sequence alignments were prepared using PROMALS3D and Jalview^58, 59^. Structure interfaces were analysed using COCOMAPS^60^.

## Description of Supplementary Movies

File name: Supplementary_Movie1.mp4

Description: **Animated 3D rendering of the family of structures from 3D variability analysis (3DVA) in CryoSPARC.** 3DVA was carried out with a dimer of HMPV N-RNA and the first variability component showed a prominent transverse tilting motion of N protomers, visualized here as a movie of related maps.

File name: Supplementary_Movie2.mp4

Description: **Simulated trajectory of a HMPV N-RNA/P 5-mer**. Snapshots from the MD simulation were taken every nanosecond. N is colored in blue (NTD in dark blue, CTD in light blue) and RNA is colored in yellow. P_CT_ peptides are colored in light red, with one peptide (colored in dark red) unbinding from a N protomer and rebinding at a neighboring protomer in the course of the simulation.

## Data Availability

Data are available on request from the corresponding authors and/or are included in the publication. Cryo-EM density maps and corresponding structure models are deposited in the Electron Microscopy Databank and the Protein Data Bank with the following accession codes: HMPV N-RNA 10mer (PDB ID: XXXX, EMD-XXXXX), HMPV N-RNA 11mer (PDB ID: XXXX, EMD-XXXXX), HMPV N-RNA spiral (PDB ID: XXXX, EMD-XXXXX), local refinement of a HMPV N-RNA dimer (PDB ID: XXXX, EMD-XXXXX), HMPV N-RNA/P 10mer (PDB ID: XXXX, EMD-XXXXX), HMPV N-RNA/P 11mer (PDB ID: XXXX, EMD-XXXXX), HMPV N-RNA/P spiral (PDB ID: XXXX, EMD-XXXXX), local refinement of a HMPV N-RNA/P dimer (PDB ID: XXXX, EMD-XXXXX). Source data are provided with the paper. Uncropped gels and Western blots are supplied in the Supplementary Information.

## Acknowledgements

We are grateful to B. van den Hoogen (Erasmus MC, Rotterdam) for providing plasmids of the HMPV NL99-1 strain. JDW is supported by a Wellcome Trust DPhil scholarship. Electron microscopy was conducted at the OPIC electron microscopy facility, which was funded by a Wellcome JIF award (060208/Z/00/Z) and was supported by a Wellcome equipment grant (093305/Z/10/Z). The Wellcome Trust is also acknowledged for providing administrative support (Grant 075491/Z/04). The work also received financial support from the French Agence Nationale de la Recherche, specific programs ANR DecRisP n° ANR-19-CE11-0017. Computation used the Oxford Biomedical Research Computing (BMRC) facility, a joint development between the Wellcome Centre for Human Genetics and the Big Data Institute, supported by Health Data Research UK and the NIHR Oxford Biomedical Research Centre. Financial support was provided by a Wellcome Trust Core Award (203141/Z/16/Z). The views expressed are those of the author(s) and not necessarily those of the NHS, the NIHR, or the Department of Health.

## Contributions

J.D.W., C.L., J.F.E., M.G., and M.R. conceived and designed the study. J.D.W. and M.R. prepared the samples for structural analysis. J.D.W, L.C., and M.R. collected cryo-EM data, processed the data, and refined the structural models. H.D., J.F., J.F.E., and M.G. performed functional assays and analyzed data. C.L. and M.R. prepared and performed MD simulations and analyzed the trajectories. J.D.W., M.G., and M.R. wrote the paper with input from all the co-authors. M.G. and M.R. supervised the work.

## References

1. van den Hoogen, B. G., et al. A newly discovered human pneumovirus isolated from young children with respiratory tract disease. Nat Med 7, (2001).

2. Jesse, S. T., Ludlow, M. & Osterhaus, A. D. M. E. Zoonotic Origins of Human Metapneumovirus: A Journey from Birds to Humans. Viruses 14, (2022).

3. Williams, J. V et al. Human metapneumovirus and lower respiratory tract disease in otherwise healthy infants and children. N Engl J Med 350, 443–50 (2004).

4. Schildgen, V. et al. Human Metapneumovirus: lessons learned over the first decade. Clin Microbiol Rev 24, 734–54 (2011).

5. Jartti, T., van den Hoogen, B., Garofalo, R. P., Osterhaus, A. D. M. E. & Ruuskanen, O. Metapneumovirus and acute wheezing in children. Lancet 360, 1393–4 (2002).

6. Midgley, C. M. et al. Notes from the Field: Severe Human Metapneumovirus Infections - North Dakota, 2016. MMWR Morb Mortal Wkly Rep 66, 486–488 (2017).

7. Edwards, K. M. et al. Burden of human metapneumovirus infection in young children. N Engl J Med 368, 633–43 (2013).

8. Haas, L. E. M., Thijsen, S. F. T., van Elden, L. & Heemstra, K. A. Human metapneumovirus in adults. Viruses 5, 87–110 (2013).

9. Afonso, C. L. et al. Taxonomy of the order Mononegavirales: update 2016. Arch Virol 161, 2351–60 (2016).

10. van den Hoogen, B. G., Bestebroer, T. M., Osterhaus, A. D. M. E. & Fouchier, R. A. M. Analysis of the genomic sequence of a human metapneumovirus. Virology 295, 119– 32 (2002).

11. Masante, C. et al. The human metapneumovirus small hydrophobic protein has properties consistent with those of a viroporin and can modulate viral fusogenic activity. J Virol 88, 6423–33 (2014).

12. Leyrat, C., Paesen, G. C., Charleston, J., Renner, M. & Grimes, J. M. Structural insights into the human metapneumovirus glycoprotein ectodomain. J Virol 88, (2014).

13. Más, V. et al. Engineering, Structure and Immunogenicity of the Human Metapneumovirus F Protein in the Postfusion Conformation. PLoS Pathog 12, e1005859 (2016).

14. Bermingham, A. & Collins, P. L. The M2– protein of human respiratory syncytial virus is a regulatory factor involved in the balance between RNA replication and transcription. Proceedings of the National Academy of Sciences 96, 11259–11264 (1999).

15. Leyrat, C., Renner, M., Harlos, K., Huiskonen, J. T. & Grimes, J. M. Drastic changes in conformational dynamics of the antiterminator M2-1 regulate transcription efficiency in pneumovirinae. Elife 2014, (2014).

16. Leyrat, C., Renner, M., Harlos, K., Huiskonen, J. T. & Grimes, J. M. Structure and self-assembly of the calcium binding matrix protein of human metapneumovirus. Structure 22, (2014).

17. Fearns, R. The Respiratory Syncytial Virus Polymerase: A Multitasking Machine. Trends Microbiol 27, 969–971 (2019).

18. Tawar, R. G. et al. Crystal structure of a nucleocapsid-like nucleoprotein-RNA complex of respiratory syncytial virus. Science 326, 1279–83 (2009).

19. Gonnin, L., et al. Structural landscape of the Respiratory Syncytial Virus nucleocapsids. bioRxiv (2023) doi:10.1101/2023.02.14.528440.

20. Pan, J. et al. Structure of the human metapneumovirus polymerase phosphoprotein complex. Nature 577, (2020).

21. Lopez, N. et al. Deconstructing virus condensation. PLoS Pathog 17, e1009926 (2021).

22. Rincheval, V. et al. Functional organization of cytoplasmic inclusion bodies in cells infected by respiratory syncytial virus. Nat Commun 8, 563 (2017).

23. Cifuentes-Muñoz, N., Branttie, J., Slaughter, K. B. & Dutch, R. E. Human Metapneumovirus Induces Formation of Inclusion Bodies for Efficient Genome Replication and Transcription. J Virol 91, (2017).

24. Boggs, K. B. et al. Human Metapneumovirus Phosphoprotein Independently Drives Phase Separation and Recruits Nucleoprotein to Liquid-Like Bodies. mBio 13, e0109922 (2022).

25. Galloux, M. et al. Minimal Elements Required for the Formation of Respiratory Syncytial Virus Cytoplasmic Inclusion Bodies In Vivo and In Vitro. mBio 11, (2020).

26. Risso-Ballester, J. et al. A condensate-hardening drug blocks RSV replication in vivo. Nature 595, 596–599 (2021).

27. Thompson, R. E., Edmonds, K. & Dutch, R. E. Specific Residues in the C-Terminal Domain of the Human Metapneumovirus Phosphoprotein Are Indispensable for Formation of Viral Replication Centers and Regulation of the Function of the Viral Polymerase Complex. J Virol e0003023 (2023) doi:10.1128/jvi.00030-23.

28. Decool, H. et al. Characterization of the Interaction Domains between the Phosphoprotein and the Nucleoprotein of Human Metapneumovirus. J Virol 96, (2022).

29. Renner, M. et al. Nucleocapsid assembly in pneumoviruses is regulated by conformational switching of the N protein. Elife 5, e12627 (2016).

30. Leyrat, C., Renner, M., Harlos, K. & Grimes, J. M. Solution and crystallographic structures of the central region of the phosphoprotein from human metapneumovirus. PLoS One 8, (2013).

31. Renner, M. et al. Structural dissection of human metapneumovirus phosphoprotein using small angle x-ray scattering. Sci Rep 7, (2017).

32. Ouizougun-Oubari, M. et al. A Druggable Pocket at the Nucleocapsid/Phosphoprotein Interaction Site of Human Respiratory Syncytial Virus. J Virol 89, 11129–43 (2015).

33. Bajorek, M. et al. Tetramerization of Phosphoprotein is Essential for Respiratory Syncytial Virus Budding while its N Terminal Region Mediates Direct Interactions with the Matrix Protein. J Virol 95, (2021).

34. Vijayakrishnan, S., et al. Ultrastructural characterization of a viral RNA and G-protein containing, membranous organelle formed in respiratory syncytial virus infected cells. bioRxiv (2022) doi:10.1101/2022.11.28.517999.

35. Conley, M. J. et al. Helical ordering of envelope-associated proteins and glycoproteins in respiratory syncytial virus. EMBO J 41, e109728 (2022).

36. Punjani, A. & Fleet, D. J. 3D variability analysis: Resolving continuous flexibility and discrete heterogeneity from single particle cryo-EM. J Struct Biol 213, 107702 (2021).

37. Khodjoyan, S. et al. Investigation of the Fuzzy Complex between RSV Nucleoprotein and Phosphoprotein to Optimize an Inhibition Assay by Fluorescence Polarization. Int J Mol Sci 24, (2022).

38. Emenecker, R. J., Griffith, D. & Holehouse, A. S. Metapredict: a fast, accurate, and easy-to-use predictor of consensus disorder and structure. Biophys J 120, 4312–4319 (2021).

39. Gilman, M. S. A. et al. Structure of the Respiratory Syncytial Virus Polymerase Complex. Cell 179, (2019).

40. Cao, D. et al. Cryo-EM structure of the respiratory syncytial virus RNA polymerase. Nat Commun 11, (2020).

41. Babu, M. M. The contribution of intrinsically disordered regions to protein function, cellular complexity, and human disease. Biochem Soc Trans 44, 1185–1200 (2016).

42. Pereira, N. et al. New Insights into Structural Disorder in Human Respiratory Syncytial Virus Phosphoprotein and Implications for Binding of Protein Partners. J Biol Chem 292, 2120–2131 (2017).

43. Choi, K. H. Viral polymerases. Adv Exp Med Biol 726, 267–304 (2012).

44. Curran, J. A role for the Sendai virus P protein trimer in RNA synthesis. J Virol 72, 4274–80 (1998).

45. Rhodin, M. H. J. et al. EDP-938, a novel nucleoprotein inhibitor of respiratory syncytial virus, demonstrates potent antiviral activities in vitro and in a non-human primate model. PLoS Pathog 17, e1009428 (2021).

46. Ahmad, A. et al. EDP-938, a Respiratory Syncytial Virus Inhibitor, in a Human Virus Challenge. N Engl J Med 386, 655–666 (2022).

47. Buchholz, U. J., Finke, S. & Conzelmann, K. K. Generation of bovine respiratory syncytial virus (BRSV) from cDNA: BRSV NS2 is not essential for virus replication in tissue culture, and the human RSV leader region acts as a functional BRSV genome promoter. J Virol 73, 251–9 (1999).

48. Berrow, N. S. et al. A versatile ligation-independent cloning method suitable for high-throughput expression screening applications. Nucleic Acids Res 35, e45 (2007).

49. Punjani, A., Rubinstein, J. L., Fleet, D. J. & Brubaker, M. A. CryoSPARC: Algorithms for rapid unsupervised cryo-EM structure determination. Nat Methods 14, 290–296 (2017).

50. Casañal, A., Lohkamp, B. & Emsley, P. Current developments in Coot for macromolecular model building of Electron Cryo-microscopy and Crystallographic Data. Protein Science 29, 1069–1078 (2020).

51. Liebschner, D. et al. Macromolecular structure determination using X-rays, neutrons and electrons: Recent developments in Phenix. Acta Crystallogr D Struct Biol 75, 861– 877 (2019).

52. Jo, S., Kim, T., Iyer, V. G. & Im, W. CHARMM-GUI: a web-based graphical user interface for CHARMM. J Comput Chem 29, 1859–65 (2008).

53. Abraham, M. J. et al. GROMACS: High performance molecular simulations through multi-level parallelism from laptops to supercomputers. SoftwareX 1–2, 19–25 (2015).

54. Lee, J. et al. CHARMM-GUI Input Generator for NAMD, GROMACS, AMBER, OpenMM, and CHARMM/OpenMM Simulations Using the CHARMM36 Additive Force Field. J Chem Theory Comput 12, 405–13 (2016).

55. Huang, J. et al. CHARMM36m: an improved force field for folded and intrinsically disordered proteins. Nat Methods 14, 71–73 (2017).

56. Pettersen, E. F. et al. UCSF Chimera--a visualization system for exploratory research and analysis. J Comput Chem 25, 1605–12 (2004).

57. Pettersen, E. F. et al. UCSF ChimeraX: Structure visualization for researchers, educators, and developers. Protein Sci 30, 70–82 (2021).

58. Waterhouse, A. M., Procter, J. B., Martin, D. M. A., Clamp, M. & Barton, G. J. Jalview Version 2--a multiple sequence alignment editor and analysis workbench. Bioinformatics 25, 1189–1191 (2009).

59. Pei, J., Kim, B.-H. & Grishin, N. V. PROMALS3D: a tool for multiple protein sequence and structure alignments. Nucleic Acids Res 36, 2295–300 (2008).

60. Vangone, A., Spinelli, R., Scarano, V., Cavallo, L. & Oliva, R. COCOMAPS: a web application to analyze and visualize contacts at the interface of biomolecular complexes. Bioinformatics 27, (2011).

