## Supplementary Material for "Structure of the N-RNA/P interface reveals mode of L/P attachment to the nucleocapsid of human metapneumovirus"

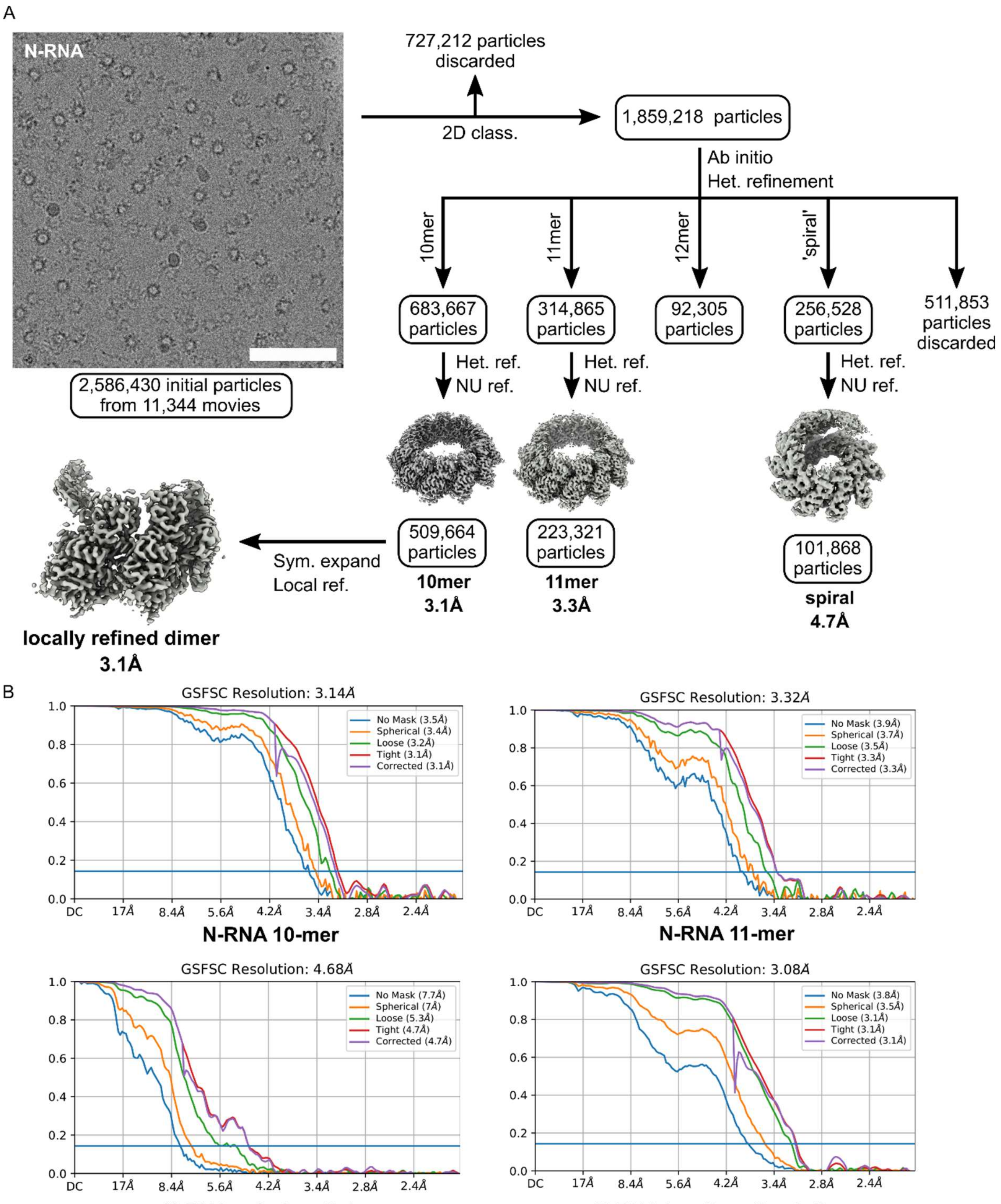


**Supplementary Fig. 1: Processing of cryo-EM data of HMPV N-RNA.** (A) Processing scheme of the HMPV N-RNA dataset in CryoSPARC. A representative image of the sample is shown on the left. Scale bar: 75 nm. (B) Fourier shell correlation (FSC) plots of the corresponding reconstructions shown in A.

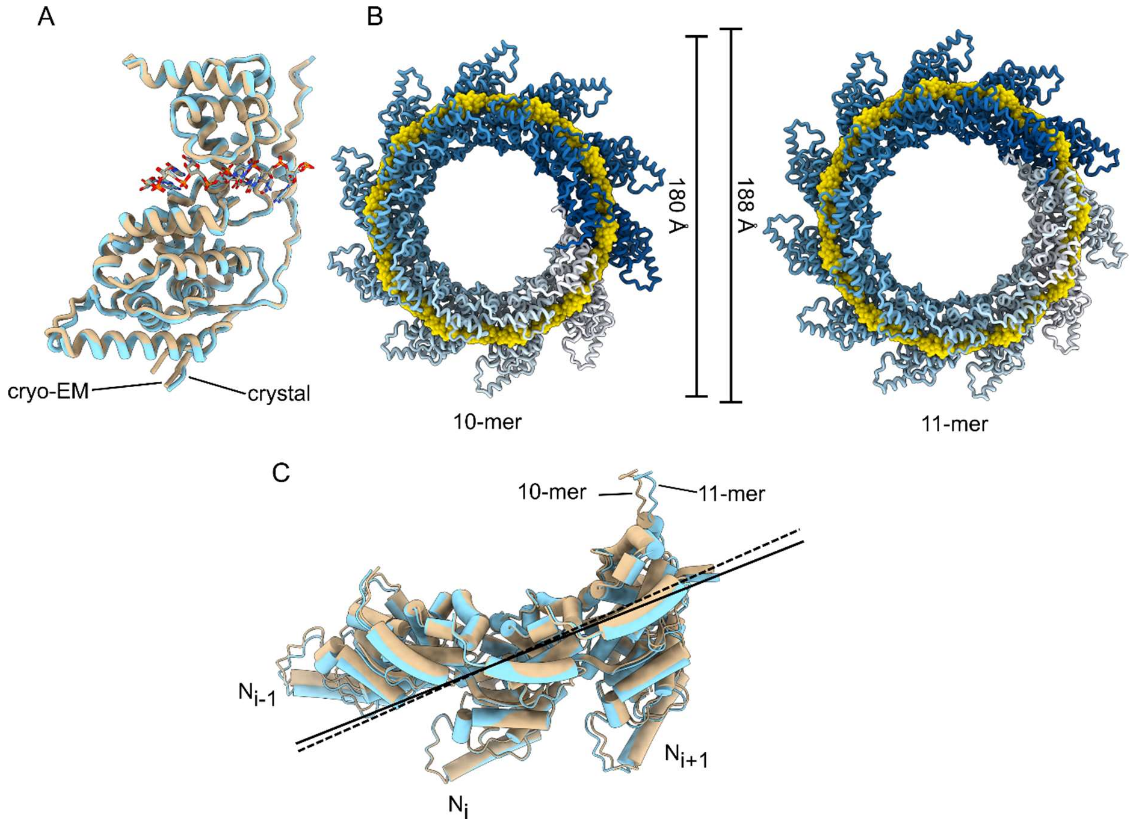


**Supplementary Fig. 2: Comparison of N-RNA oligomers.** (A) Structural alignment of N-RNA protomers from cryo-EM (this study, beige) and X-ray crystallography (PDB ID: 5FVC, light blue). (B) Comparison of diameter of N-RNA 10-mers and 11-mers (indicated). (C) Structural alignment of three neighboring N-RNA protomers from a 10-mer (beige) and 11-mer (light blue). The alignment shows a slightly different angle between neighboring N-protomers from 10-mers and 11-mers, required to accommodate the ring expansion.

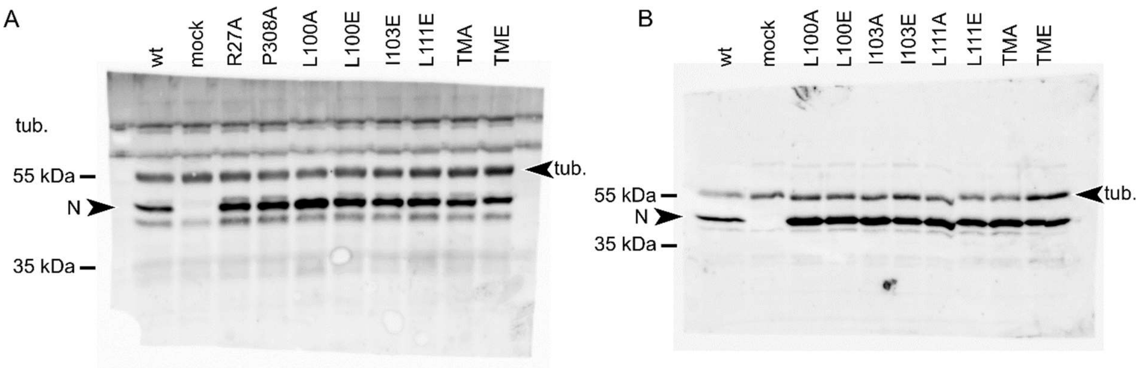


**Supplementary Fig. 3: Uncropped Western blots.** (A - B) Validation by Western blot of N expression (wt and mutants) in BSRT-7 cells transfected in the context of minigenome assays. The height of N bands is indicated.

A

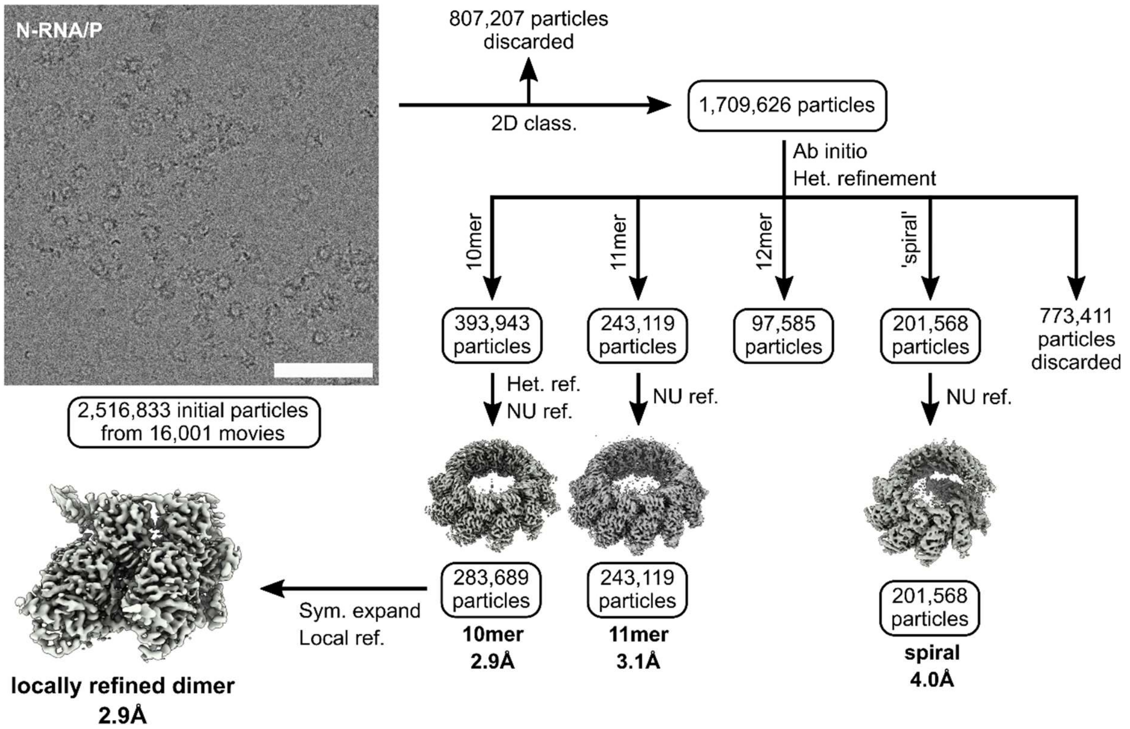


B

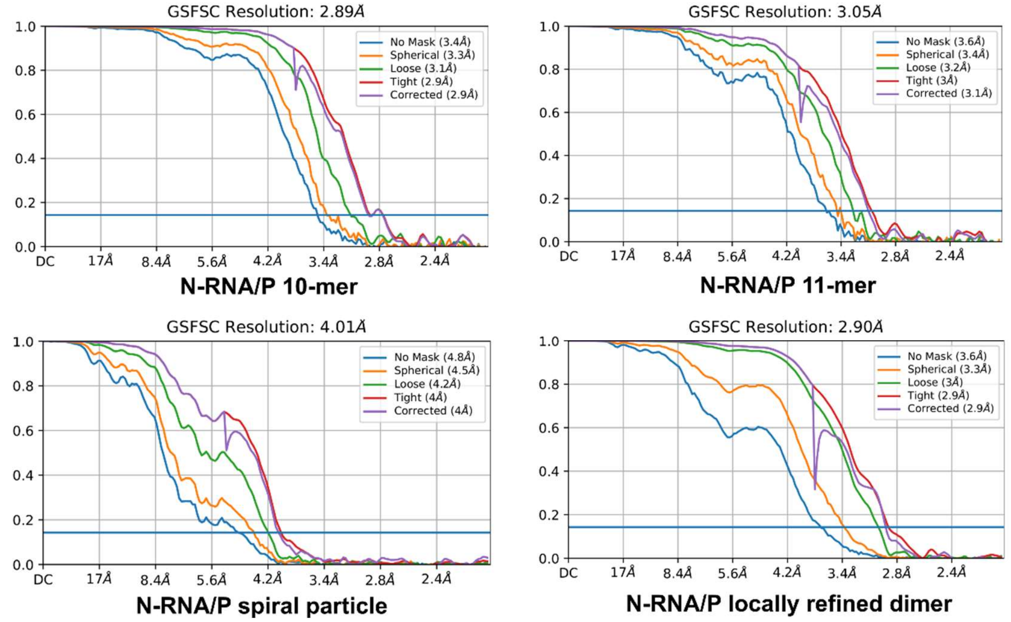


**Supplementary Fig. 4: Processing of cryo-EM data of HMPV N-RNA incubated with $P_{CT}$ peptide.** (A) Processing scheme of the HMPV N-RNA/P dataset in CryoSPARC. A representative image of the sample is shown on the left. Scale bar: 75 nm. (B) Fourier shell correlation (FSC) plots of the corresponding reconstructions shown in A.

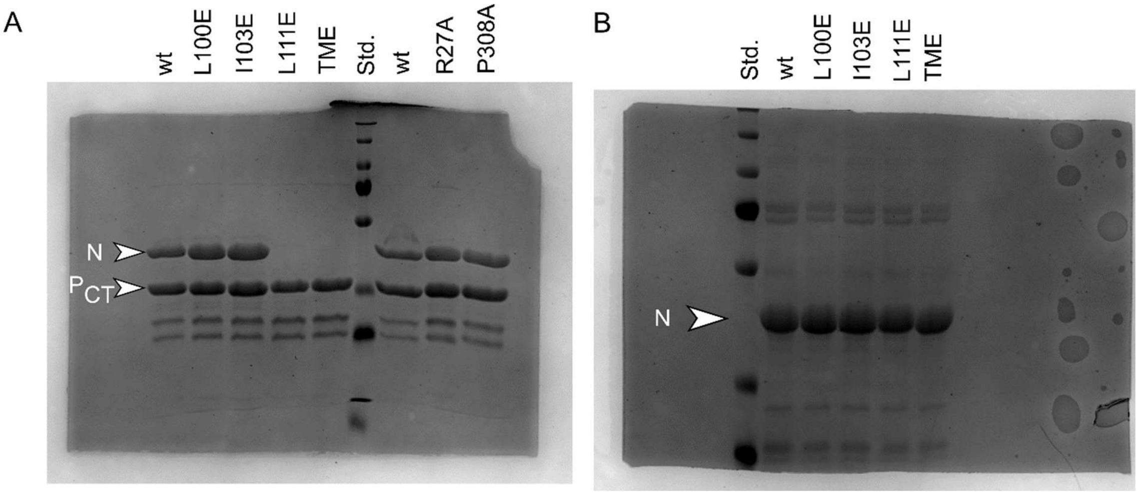


**Supplementary Fig. 5: Uncropped SDS-PAGE gels.** (A) GST-PCT and 6xHis-tagged N proteins (wt and mutants) were co-expressed and co-purified using a GST-tag. A SDS-PAGE analysis of the products of purification is shown, with the position of N and GST-$P_{CT}$ indicated. (B) SDS-PAGE analysis of N mutants (as indicated) purified using 6xHis-tag.

| Dataset | HMPV N-RNA dataset | | | | HMPV N-RNA/P dataset | | | |
|---|---|---|---|---|---|---|---|---|
| **Structure** | **N-RNA 10-mer** | **N-RNA 11-mer** | **N-RNA Spiral*** | **N-RNA dimer Loc. ref.** | **N-RNA/P 10-mer** | **N-RNA/P 11-mer** | **N-RNA/P Spiral*** | **N-RNA/P dimer Loc. ref** |
| **Data collection** | | | | | | | | |
| Microscope | Titan Krios | | | | Titan Krios | | | |
| Voltage (kV) | 300 | | | | 300 | | | |
| Detector | Gatan K2 | | | | Gatan K2 | | | |
| Magnification | 130,000x | | | | 130,000x | | | |
| Pixel size (Å) | 1.05 | | | | 1.05 | | | |
| Total dose ($e^{-}/Å^{2}$) | 44.2 | | | | 44.2 | | | |
| Def. range (µm) | -1.0 to -2.6 | | | | -1.0 to -2.6 | | | |
| Frames/movie | 50 | | | | 50 | | | |
| **Data processing** | | | | | | | | |
| Number of movies | 11,344 | | | | 16,001 | | | |
| Initial particles | 2,586,430 | | | | 2,516,833 | | | |
| Box size (pixels) | 400 | | | | 400 | | | |
| Final particles | 509,664 | 223,321 | 101,868 | 2,548,320 sym. exp.# | 283,689 | 243,119 | 201,568 | 1,418,445 sym. exp.# |
| Resolution (FSC 0.143) | 3.1 Å | 3.3 Å | 4.7 Å | 3.1 Å | 2.9 Å | 3.1 Å | 4.0 Å | 2.9 Å |
| Symmetry | C10 | C11 | C1 | C1 | C10 | C11 | C1 | C1 |
| Sharp. B-factor ($Å^{2}$) | -133 | -131 | -138 | -87 | -124 | -121 | -108 | -70 |
| **Model Refinement** | | | | | | | | |
| Composition | | | | | | | | |
| Protein residues | 351 | 351 | - | 702 | 370 | 370 | - | 744 |
| Nucleotides | 7 | 7 | - | 14 | 7 | 7 | - | 14 |
| Atoms | 2861 | 2861 | - | 5722 | 3016 | 3016 | - | 6056 |
| ADP | | | | | | | | |
| Protein | 46.39 | 49.92 | - | 32.41 | 39.08 | 33.95 | - | 30.52 |
| Nucleotides | 26.63 | 38.20 | - | 23.51 | 24.21 | 21.49 | - | 17.52 |
| RMSD from ideal | | | | | | | | |
| Bond lengths (Å) | 0.008 | 0.006 | - | 0.006 | 0.004 | 0.004 | - | 0.004 |
| Bond angles (°) | 0.820 | 0.722 | - | 0.652 | 0.570 | 0.553 | - | 0.568 |
| Validation | | | | | | | | |
| MolProbity score | 1.47 | 1.52 | - | 1.45 | 1.46 | 1.45 | - | 1.43 |
| Clashscore | 8.75 | 9.80 | - | 8.31 | 8.61 | 8.28 | - | 7.83 |
| Rotamer outliers (%) | 0.00 | 0.00 | - | 0.00 | 0.00 | 0.00 | - | 0.00 |
| CC model vs. map | 0.79 | 0.78 | - | 0.81 | 0.80 | 0.81 | - | 0.83 |
| Ramachandran | | | | | | | | |
| Favored (%) | 98.27 | 97.98 | - | 98.13 | 98.63 | 98.08 | - | 98.37 |
| Allowed (%) | 1.73 | 2.02 | - | 1.87 | 1.37 | 1.92 | - | 1.63 |
| Outliers (%) | 0.00 | 0.00 | - | 0.00 | 0.00 | 0.00 | - | 0.00 |
| Data Deposition | | | | | | | | |
| PDB | XXXX | XXXX | XXXX | XXXX | XXXX | XXXX | XXXX | XXXX |
| EMDB | XXXX | XXXX | XXXX | XXXX | XXXX | XXXX | XXXX | XXXX |

**Supplementary Table 1: Cryo-EM data collection and refinement statistics.** Only a single asymmetric unit (ASU) was refined when C10 or C11 symmetry was applied.
*For spiral particles, atomic refinement was not carried out (only rigid-body) due to insufficient map quality.
#Number of particles following from 5-fold symmetry expansion
